# Structural mechanism defining product specificity in glycoside hydrolase family 66 cycloisomaltotetraose glucanotransferase

**DOI:** 10.64898/2026.08.30.748175

**Authors:** Ryota Yasukochi, Toma Kashima, Tetsuya Mori, Yuki Kawauchi, Akimasa Miyanaga, Hikaru Watanabe, Shinya Fushinobu

## Abstract

Cyclic oligosaccharides possess industrial advantages, including molecular encapsulation capability and high physicochemical stability, owing to the absence of a reducing end. Recently, a novel cyclic tetrasaccharide, cycloisomaltotetraose (CI4), consisting of four α-1,6-linked glucose units, and the enzymes responsible for its synthesis, cycloisomaltotetraose glucanotransferases (CI4Tases), were discovered. Unlike known cycloisomaltooligosaccharide glucanotransferases (CITases) that yield a wide distribution of cyclic products with a degree of polymerization (DP) of 7 or higher, CI4Tases strictly produce CI4. To elucidate the molecular mechanism underlying this strict DP4 specificity, we determined the crystal structures of CI4Tase from *Agreia* sp. D1110, in its ligand-free form, as well as in complex with the linear hydrolysis product isomaltotetraose (IG4) and with CI4. Structural comparisons revealed that a loop (M247 to R251) blocks the region corresponding to the –5 subsite of typical CITases, narrowing the substrate-binding pocket. This “molecular ruler” mechanism ensures that only a glycan chain of exactly four glucose units is accommodated for cyclization. Among mutants of the residue positioned at the center of bound CI4, the formation of by-products other than CI4 was significantly suppressed in F245L, F245A, and F245W. While the cyclization activity of all F245 mutants decreased, the CI4 hydrolysis activity of these three mutants was also significantly reduced, resulting in an increased specificity for cyclic sugar production. These findings elucidate the strict size-control mechanism of CI4Tase and provide a structural foundation for engineering cycloisomaltooligosaccharide-producing enzymes with optimized transglycosylation efficiency and specificity for industrial applications.

## Introduction

Cyclic oligosaccharides are ring-structured carbohydrates lacking a reducing end, a feature that protects them from Maillard reactions and confers exceptional thermal and pH stability. Their hydrophobic central cavities can form inclusion complexes with a wide range of guest molecules, making them highly valuable for applications in food, cosmetics, pharmaceuticals, and environmental technology [1,2]. Among these, cyclodextrins with degrees of polymerization (DP) ranging from 6 to 8 (α-, β-, and γ-CD with α-1,4 linkages) are the most well-known cyclic sugars, typically synthesized from starch by cyclodextrin glucanotransferase (CGTase, EC 2.4.1.19) [3]. In addition to CDs, other starch-derived cyclic tetrasaccharides, such as cyclic nigerosylnigerose (CNN or cycloalternan) [4–6], and cyclic maltosyl-maltose (CMM) [7–9], have been identified. Among α-1,6-linked cyclic glucans, cyclic isomaltooligosaccharides (CIs) typically exhibit DPs ranging from 7 to 10 or higher [10]. CIs not only have higher water solubility and molecular encapsulation capability than CDs [11], but they also exhibit anti-cariogenic activity by inhibiting glucosyltransferases from *Streptococcus mutans* [12,13]. CIs are synthesized from dextran by cycloisomaltooligosaccharide glucanotransferases (CITases, EC 2.4.1.248) [14], which belong to the glycoside hydrolase (GH) family 66 in the Carbohydrate-Active enZymes (CAZy) database [15]. To date, CITases from *Bacillus circulans* (currently reclassified as *Paenibacillus agaridevorans*) T-3040 (BcCITase) [16] and *Paenibacillus* sp. 598K (PsCITase) [17] have been structurally and functionally characterized. While the major DP produced by CITases varies depending on the specific enzyme (e.g., BcCITase mainly produces CI8, whereas PsCITase mainly produces CI7), a single enzymatic reaction generally yields a diverse profile of cyclic products, typically ranging from DP7 to DP17, and in some cases exceeding DP20 [18].

Recently, we identified a novel cyclic tetrasaccharide termed cycloisomaltotetraose (CI4; *cyclo*-{→6)-α-D-Glc*p*-(1→6)-α-D-Glc*p*-(1→6)-α-D-Glc*p*-(1→6)-α-D-Glc*p*-(1→}) [19] and its synthesizing enzymes from two soil bacterial strains [20]. These enzymes catalyze four reactions: (a) cyclization of isomaltooligosaccharides (IGs, DP > 5) into CI4, (b) disproportionation, (c) hydrolysis of CI4, and (d) coupling reactions between CI4 and IGs (Fig. 1). The CI4-forming cyclization is the main reaction, whereas the others are minor side reactions. Notably, although a low level of hydrolytic activity toward cyclic CI4 was detected, these enzymes exhibit almost no hydrolytic activity toward linear IGs. Unlike conventional CITases, these newly discovered enzymes generate CI4 as the sole cyclic product. Based on this strict size specificity, the enzymes were designated as cycloisomaltotetraose glucanotransferase (CI4Tase, EC 2.4.1.–). The CI4Tase gene products identified from the genome sequences of *Agreia* sp. D1110 and *Microbacterium trichothecenolyticum* D2006, originally designated as ORF9038 and ORF5328 [21], are herein referred to as AgCI4Tase and MtCI4Tase, respectively. AgCI4Tase and MtCI4Tase share 71.3% sequence identity and exhibit approximately 37% identity with BcCITase. Both CI4Tases contain an N-terminal GH66 domain interrupted by the insertion of a carbohydrate-binding module family 35 (CBM35) domain, followed by a C-terminal CBM13 domain (Fig. 2A). Despite their potential for industrial applications, the molecular mechanism underlying how CI4Tases strictly produce a DP4 ring without forming larger CIs has remained unclear. In this study, we performed structural analysis of AgCI4Tase. Additionally, we conducted site-directed mutagenesis of a central cavity residue to investigate its effects on enzyme activity and side-reaction suppression.

**Fig. 1.**
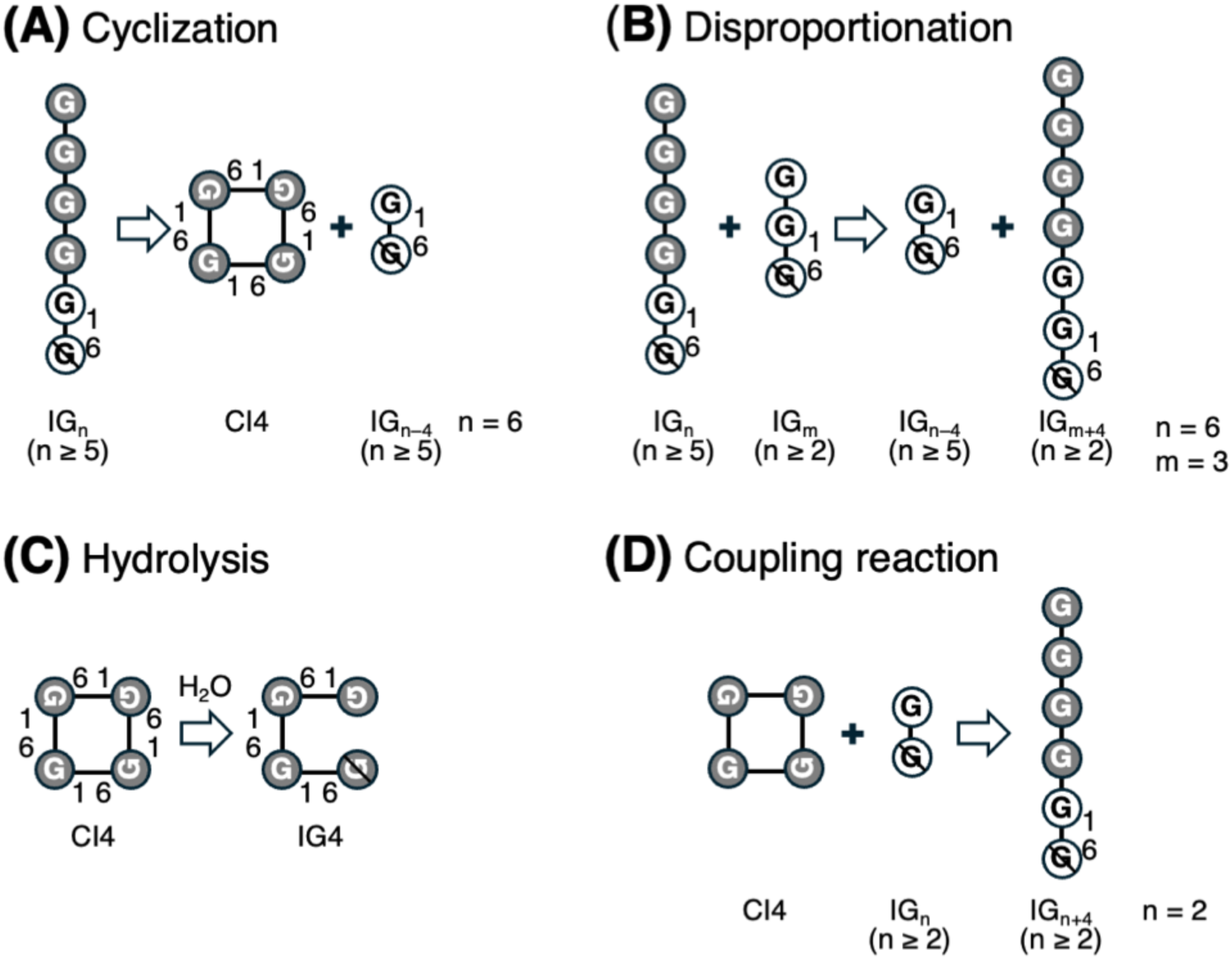
Four reactions catalyzed by CI4Tase. IG_n_, isomaltooligosaccharides with degrees of polymerization (DP) = n; CI4, cycloisomaltotetraose; IG4, isomaltotetraose. (A) CI4-producing intramolecular transglycosylation (cyclization) from IG_n_. A case of n = 6 is shown. (B) Intermolecular transglycosylation (disproportionation) of an IG4 unit to produce IG_n_ with different DPs from the substrates. A case of n = 6 and m = 3 is shown. (C) Hydrolysis reaction cleaving an α-1,6 linkage in CI4 to produce IG4. (D) Intermolecular transglycosylation (coupling reaction) of CI4 and IG_n_ to produce IG_n+4_. A case of n = 2 is shown. Reducing end glucose units are marked by a diagonal line. Adapted from Figure 7 of Fujita et al. [20].

**Fig. 2.**
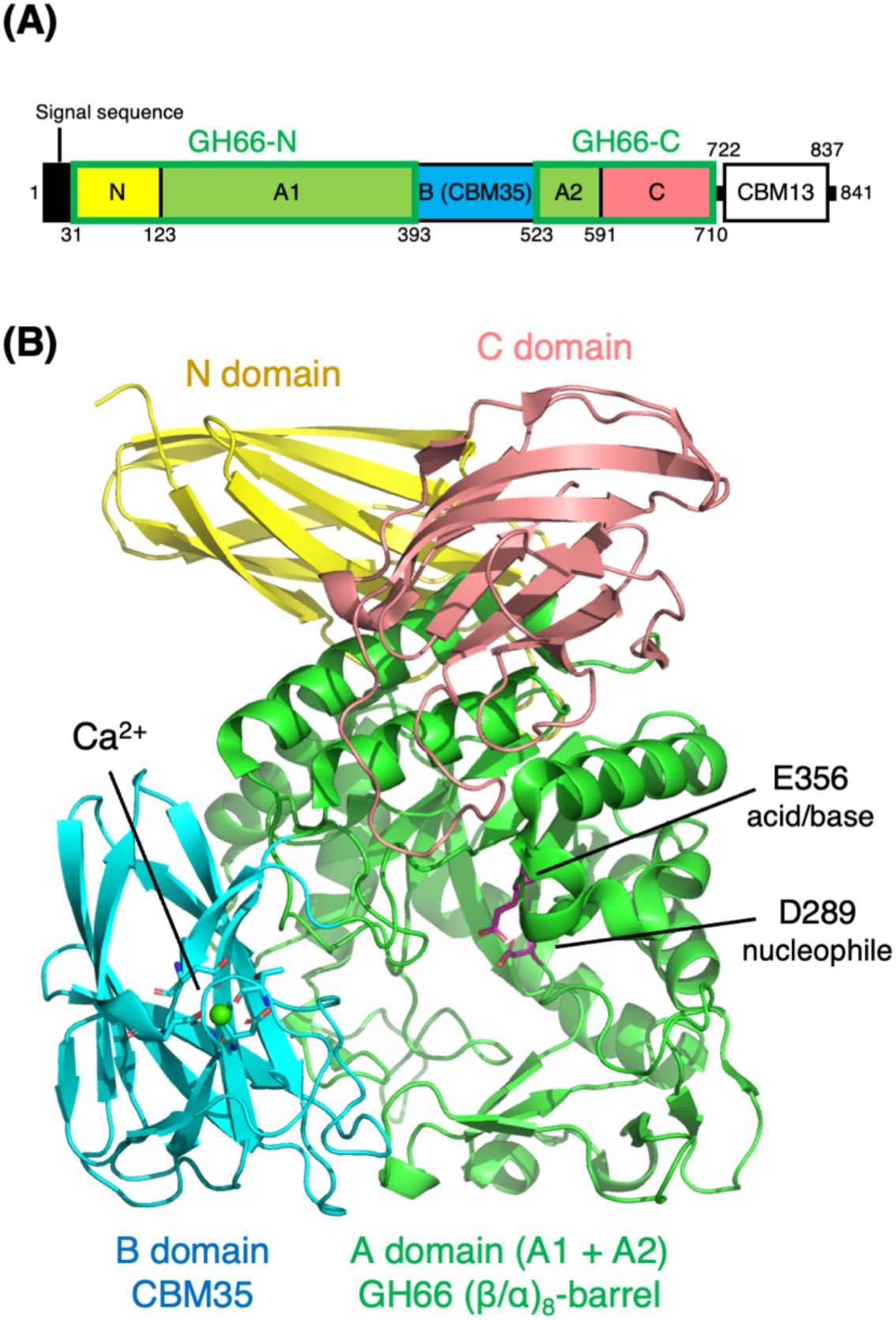
Domain architecture and crystal structure of AgCI4Tase. (A) Schematic representation of domain organization. A protein construct lacking the signal peptide and CBM13 domain was used for crystallization. (B) The overall structure of ligand-free AgCI4Tase. The structure is colored as follows: Domain N (yellow, immunoglobulin-like β-sandwich), Domain A (green, (α/β)_8_-barrel), Domain B (blue, β-jellyroll), and Domain C (salmon, Greek key β-sandwich). The catalytic residues are shown as magenta sticks.

## Results

### Overall structure

To optimize crystallization, we utilized a construct lacking the N-terminal signal sequence and the C-terminal CBM13 domain (residues 32–710; Fig. 2A), following the strategy used for BcCITase [16]. Short sequences of the SKIK tetrapeptide [22] and hexahistidine tags were introduced at the N- and C-termini, respectively, to facilitate recombinant protein expression in *Escherichia coli* and metal-affinity purification. In size-exclusion chromatography, AgCI4Tase eluted at a position corresponding to a molecular mass smaller than the theoretical molecular mass of the monomer (74.5 kDa), likely due to an interaction with the agarose column matrix (Fig. 3). The crystal structure of ligand-free, wild type (WT) AgCI4Tase was determined at 2.89 Å resolution (Fig. 2B and Table 1). The asymmetric unit of the crystal contained an AgCI4Tase monomer, consistent with an analysis of the crystal-packing interface using the PISA server [23]. The structure of AgCI4Tase consists of two large domains, GH66 (N- and C-terminal regions) and CBM35, and is further divided into four subdomains: an N-terminal immunoglobulin-like β-sandwich domain (N domain), a central (β/α)_8_-barrel domain (A1 + A2 domain), an inserted β-jellyroll domain (B domain, CBM35), and a C-terminal Greek-key β-sandwich domain (C domain) (Fig. 2B). The domain architecture of AgCI4Tase is similar to that of structurally characterized GH66 CITases, and the root-mean-square deviation (RMSD) with BcCITase (PDB ID: 3WNK) was 3.2 Å for 655 Cα atoms.

**Fig. 3.**
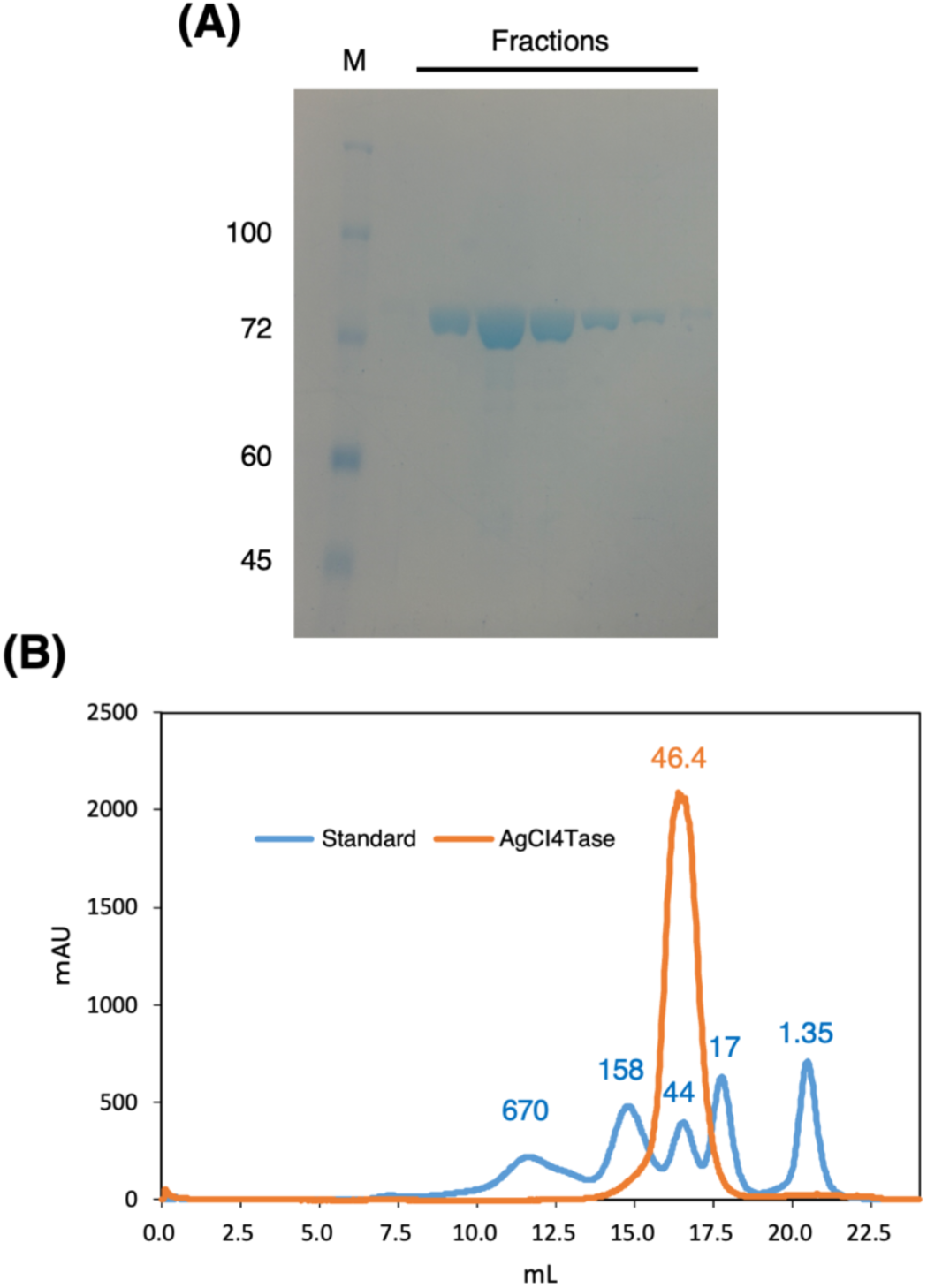
Size-exclusion chromatography. (A) SDS-PAGE analysis of the elution fractions. Lane M: molecular weight marker. (B) Size-exclusion chromatogram of AgCI4Tase (orange line) and molecular weight standards (blue line). The numbers above the peaks indicate the molecular masses (in kDa) of the standards and the estimated mass of AgCI4Tase (46.4 kDa). A Superose 6 10/300 GL column was used.

**Table 1.** Crystallographic data collection and refinement statistics of AgCI4Tase crystals.

|  | WT ligand-free | WT + IG4 | D289A + CI4 |
| --- | --- | --- | --- |
| **Data collection<sup>a</sup>** |  |  |  |
| Beamline | SPring-8 BL45XU | KEK PF AR-NE3A | SPring-8 BL45XU |
| Space group | $I4_122$ | $I4_122$ | $I4_122$ |
| Unit cell (Å) | $a = b = 216.83$,<br>$c = 169.75$ | $a = b = 214.84$,<br>$c = 168.14$ | $a = b = 217.64$,<br>$c = 166.53$ |
| Resolution (Å) | 50.23–2.89 (2.99–<br>2.89) | 49.77–2.73 (2.81–<br>2.73) | 48.67–3.01 (3.13–<br>3.01) |
| Total reflections | 2484767 (252595) | 696427 (53393) | 2198274 (249707) |
| Unique reflections | 45335 (4398) | 52173 (4458) | 39693 (4398) |
| Multiplicity | 54.8 (57.4) | 13.3 (12.0) | 55.4 (56.8) |
| Completeness (%) | 100.0 (100.0) | 100.0 (100.0) | 99.9 (99.3) |
| Mean $I/\sigma(I)$ | 13.8 (1.7) | 6.8 (1.1) | 9.1 (1.0) |
| $R_{\text{merge}}$ | 0.586 (6.697) | 0.249 (1.979) | 1.094 (9.824) |
| $R_{\text{pim}}$ | 0.591 (6.756) | 0.258 (2.068) | 1.104 (9.912) |
| $CC_{1/2}$ | 0.997 (0.683) | 0.992 (0.684) | 0.981 (0.622) |
| Wilson B-factor (Å<sup>2</sup>) | 71.40 | 60.00 | 48.40 |
| **Refinement** |  |  |  |
| Resolution range (Å) | 50.23–2.89 | 48.46–2.73 | 48.67–3.01 |
| No. of reflections | 42987 | 49558 | 37647 |
| $R_{\text{work}}/R_{\text{free}}$ | 0.190/0.229 | 0.197/0.234 | 0.209/0.244 |
| No. of atoms |  |  |  |
| Protein | 5176 | 5182 | 5180 |
| Heterogens | 1 | 94 | 45 |
| Solvent | 34 | 27 | 22 |
| Average B-factor (Å<sup>2</sup>) | 62.07 | 60.65 | 40.16 |
| RMSD from ideal values |  |  |  |
| Bond lengths (Å) | 0.0090 | 0.0079 | 0.0080 |
| Bond angles (°) | 2.196 | 1.948 | 1.981 |
| Ramachandran plot (%) |  |  |  |
| Favored | 92.22 | 93.4 | 93.4 |
| Allowed | 6.75 | 6.6 | 6.31 |
| Outlier | 1.03 | 0.00 | 0.29 |
| Clashscore | 6 | 6 | 6 |
| PDB code | 24TK | 24TL | 24TM |

<sup>a</sup>Values in parentheses represent the highest resolution shell.

A Ca²⁺ ion was observed in the CBM35 domain, which possesses a conserved Ca²⁺-binding site in GH66–CBM35 CITases [16,17]. Notably, the CBM35 domain is inserted into the loop region of the barrel fold of the A domain, a structural arrangement that is well conserved among CITases. In a previous study of BcCITase, excision of the corresponding CBM35 domain resulted in a drastic reduction in the *k*_cat_/*K*_m_ value for CI-forming activity to only 3.8% of the WT level, highlighting the functional importance of this inserted domain [24]. This type of domain insertion within the catalytic barrel domain has also been reported in other GH enzymes, including GH5 endoglucanases [25] and GH148 β-glucanases [26], where the inserted CBMs are thought to facilitate substrate recognition.

### Complex structures with hydrolysis product and cyclic tetrasaccharide

The crystal structure of WT AgCI4Tase in complex with isomaltotetraose (IG4) was determined at 2.73 Å resolution (Table 1). This complex was obtained by co-crystallization with CI4, which was hydrolyzed during crystal growth. In the active site, a linear IG4 tetrasaccharide is bound at subsites from –4 to –1 (Fig. 4A). The reducing-end glucose (α-anomer) at subsite –1 (Glc(–1)) adopts a non-catalytic orientation, with its hydroxy groups forming hydrogen bonds with the side chains of E356 (acid/base catalyst), D289 (nucleophile), and Q290. Glc(–2) is extensively recognized through hydrogen bonds involving Y161, D162, H167, and Y213. In particular, D162 forms a pair of hydrogen bonds with the O3 and O4 hydroxy groups, making a major contribution to substrate binding. In contrast, substrate recognition at subsite –3 is relatively weak, mediated by only a single hydrogen bond with the main chain of M247. The O2 and O3 hydroxy groups of Glc(–4) form hydrogen bonds with E488 and D560. Notably, E488 is the sole residue from the CBM35 domain involved in these interactions, suggesting that the direct contribution of the CBM35 domain to substrate binding is limited.

**Fig. 4.**
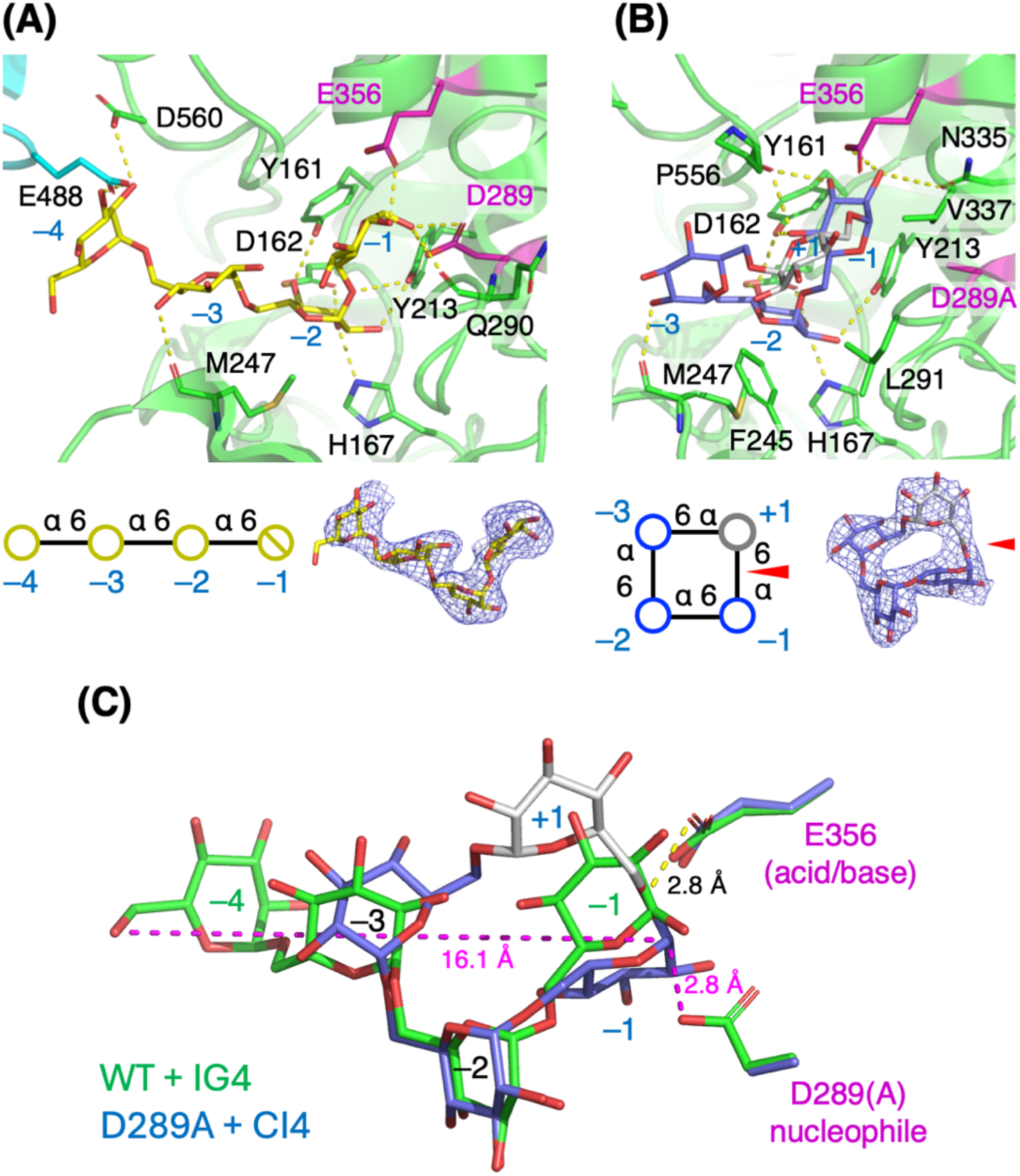
The active site structure of AgCI4Tase in complex with linear and cyclic ligands, and their superimposition. (A) WT AgCI4Tase complexed with IG4 (yellow). Residues in the CBM35 domain are shown in cyan. (B) D289A mutant complexed with CI4 (blue for glucose units in subsites –1 to – 3, and gray for the unit in subsite +1). The catalytic residues are shown as magenta. Schematic representation and Polder maps of the ligands (blue meshes contoured at 2.0σ for IG4 and 2.5σ for CI4) are shown at the bottom. The bond undergoing cleavage or transfer reaction in CI4 is indicated by a red wedge. (C) Superimposition of IG4 (green) and CI4 (the same color code as (B)) in the active site of the complex structures. Hydrogen bonds and interatomic distance are shown as dashed yellow and magenta lines.

The crystal structure of AgCI4Tase in complex with CI4 was determined at 3.01 Å resolution (Table 1). This complex was obtained by co-crystallization of the inactive nucleophile mutant (D289A) with CI4. The cyclic IG4 molecule was clearly observed in the electron density, and the binding mode of Glc(–2) is essentially identical to that observed in the IG4 complex (Fig. 4B). For Glc(–3), the hydrogen bond with the main chain of M247 is retained, whereas the sugar ring is rotated due to the restricted linkage of the small cyclic molecule. The glucose unit located between subsites –1 and –3 is assigned as Glc(+1) based on its position relative to the catalytic residues (E356 and D289A) and Glc(–1). Although Glc(+1) forms no polar interactions with the protein, it is accommodated within a hydrophobic environment formed by F245, L291, and V337. Glc(–1) in CI4 is bound at the canonical subsite –1, forming extensive hydrogen bonds with the side chains of Y161, N335, and E356, and the main chain of P556.

Superimposition of the IG4 and CI4 molecules in the complex structures revealed that Glc(– 2) is well aligned, whereas Glc(–3) and Glc(–1) deviate significantly (Fig. 4C). In the CI4 complex, the side chain of E356 (acid/base catalyst) forms a hydrogen bond with the α-1,6-glucosidic bond oxygen connecting Glc(–1) and Glc(+1). In this superimposition, the side-chain oxygen of D289 in the WT AgCI4Tase structure is positioned in-line with the scissile Glc(–1)–Glc(+1) bond of the CI4 complex structure, at a distance of 2.8 Å from the C1 atom of Glc(–1), suggesting that this residue performs the nucleophilic attack in the Michaelis complex. During the cyclization reaction, the O6 hydroxy of Glc(−4) attacks the C1 atom of Glc(−1) in the covalent intermediate. In the superimposed structures of IG4 and CI4, the distance between these atoms is 16.1 Å.

### Structural comparison with CITase

To identify the structural elements underlying the strict four-glucose-unit specificity of AgCI4Tase, we compared its active-site architecture (Fig. 5A) with that of BcCITase in complex with IG8 (Fig. 5B) [16]. While the residues at subsites –1 and –2 are highly conserved, AgCI4Tase possesses a distinct loop (M247–R251) that protrudes into the substrate-binding pocket, narrowing it relative to the more open pocket of BcCITase. Within this loop, the side chain of P249 projects directly into the path of the glucan chain, where it would sterically clash with the glucose unit occupying subsite –5 in BcCITase. This steric block caps the binding pocket, preventing the accommodation of longer isomaltooligosaccharides and thereby restricting the transglycosylation product to a strict four-glucose-unit length. In BcCITase, the deep substrate-binding pocket accommodates a longer glucan chain, forming distinct subsites beyond –5. This region is referred to as the 2^nd^ site of CBM35 [16], where Q496, Y499, and W514 interact extensively with Glc(–8) at the non-reducing end of the substrate. In contrast, AgCI4Tase forms only a single hydrogen bond with Glc(–4) via the side chain of E488, implying a significantly weaker contribution of its CBM35 domain to carbohydrate binding. In the BcCITase structure, IG4 is also bound in a cleft on the distal side of CBM35 from the active site (Fig. 5B, bottom). This binding site (1^st^ site) is presumed to play a role in binding the long-chain dextran substrate and guiding it to the active site [16]. In the structure of AgCI4Tase, glycerol added as a cryoprotectant was bound at the corresponding site (Fig. 5A, bottom).

**Fig. 5.**
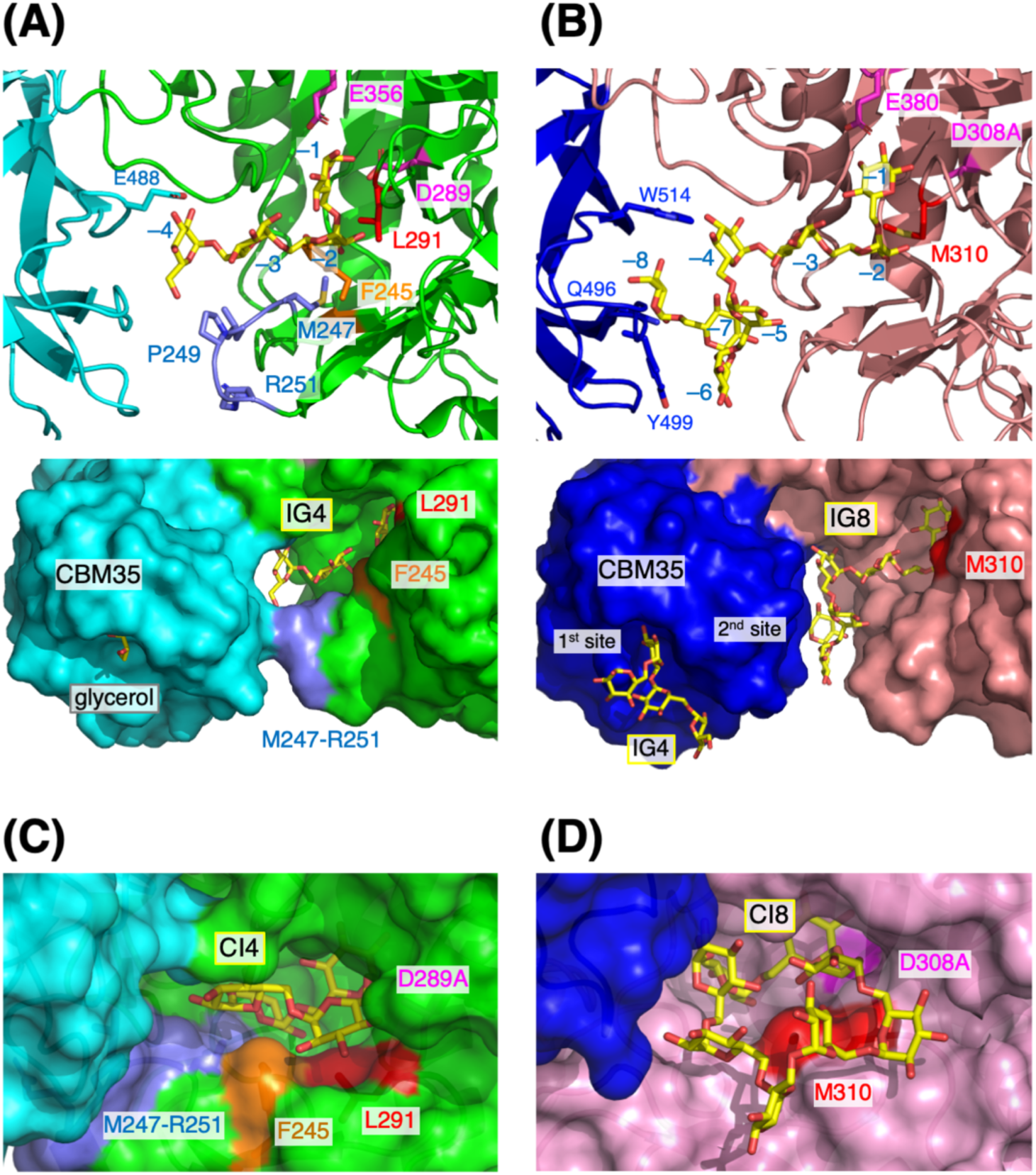
Comparison of the substrate-binding sites of AgCI4Tase and CITase from *B. circulans* (BcCITase). (A, B), cartoon-stick (top) and surface (bottom) representations of the complex structures with linear ligands. (A) AgCI4Tase (GH66, green; CBM35, cyan) complexed with IG4 (yellow sticks). The M247–R251 loop, F245 (central residue), and L291 are shown in light blue, orange, and red, respectively. The catalytic residues are shown as magenta. A glycerol molecule bound in the 1^st^ site of CBM35 is shown as gray sticks. (B) BcCITase (GH66, pink; CBM35, blue) complexed with isomaltooctaose (IG8, yellow sticks) (PDB ID: 3WNN). M310 (central residue) is shown in red. (C, D). Surface representations of the complex structures with cyclic ligands. (C) AgCI4Tase complexed with CI4. (D) BcCITase complexed with cycloisomaltooctaose (CI8) (PDB ID: 3WNO).

In cyclodextrin-producing CGTases, mutating residue Y195, positioned at the center of the cyclic sugar molecule, to another amino acid residue bearing an aromatic ring or a bulky side chain is known to affect both the product-size specificity (the ratio of α-, β-, and γ-CD) and the reaction rates [3,27]. Therefore, we compared the structures of AgCI4Tase and BcCITase in complex with their cyclic reaction products, CI4 and CI8, respectively. In BcCITase, M310, located at the center of the bound CI8 molecule (Fig. 5D), has been suggested to play a similar functional role to that of Y195 in CGTase [16]. In AgCI4Tase, the residue corresponding to M310 of BcCITase is L291 (Fig. 5C). However, the side chain of L291 is closest to Glc(+1) (minimum interatomic distance of 4.4 Å), whereas it is further distant from Glc(–2) and Glc(–3) (4.8 and 6.3 Å, respectively). In contrast, the side chain of the adjacent F245 lies close to Glc(–2) and Glc(–3) (minimum interatomic distances of 3.4 and 3.7 Å, respectively), placing it in a better position to interact with the hydrophobic center of the cyclic sugar during the cyclization reaction. Therefore, F245 in AgCI4Tase is suggested to be the functional counterpart of M310 in BcCITase.

### Mutational analysis of F245

To investigate the functional significance of residue F245, we generated the F245A, F245L, F245W, and F245Y mutants of AgCI4Tase and evaluated their catalytic activity. Time-course analysis of the enzymatic reaction using dextran as a substrate by thin-layer chromatography (TLC, Fig. 6) showed that WT mainly produced CI4 at 10 min. However, as the reaction progressed, linear oligosaccharides of various lengths were generated through side reactions, including coupling and disproportionation, and hydrolysis (Fig. 1). As with the WT enzyme, the mutant enzymes produced CI4 as the major product, along with the same by-products corresponding to the spots above and below CI4. Interestingly, the F245A, F245L, and F245W mutants showed markedly reduced accumulation of side-reaction products. In particular, F245A showed a reduction in two spots with higher retardation factor (*R*_f_) values than CI4. These spots correspond to isomaltose (IG2) and isomaltotriose (IG3), indicating that F245A suppresses the formation of short oligosaccharides via disproportionation reaction. In F245L, the production of oligosaccharides of IG5 and longer was particularly reduced, and the spot immediately below CI4, corresponding to IG4, was also barely detectable, suggesting that hydrolytic activity was suppressed. For F245Y, the production of side-reaction products was slightly increased compared with WT.

**Fig. 6.**
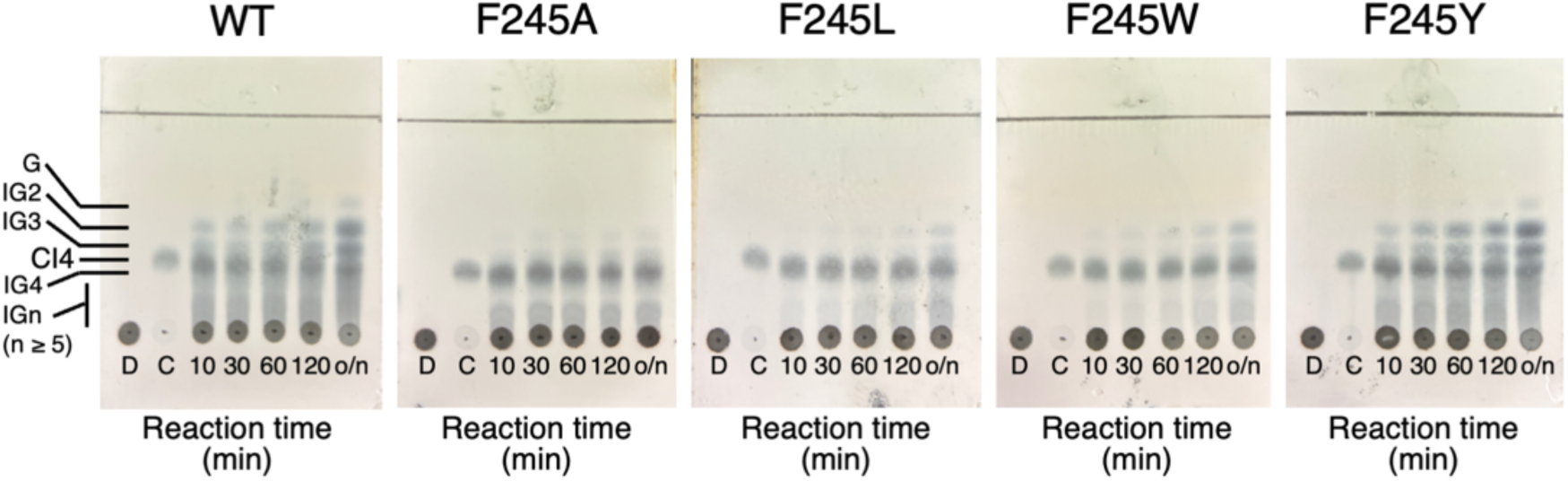
Time-course TLC analysis of CI4 production by WT AgCI4Tase and F245 mutants. Reaction mixtures containing 1% dextran, 50 mM Na-acetate (pH 6.0), and purified enzyme (0.1 mg/mL) were incubated at 37°C. G, glucose; IGn, isomaltooligosaccharides with DP = n; D, 1% dextran; C, 1 mM CI4.

The cyclization activity producing CI4 from dextran was also measured by high-performance liquid chromatography (HPLC, Fig. 7). The chromatographic profile of the reaction products generated by F245L was similar to that of the WT enzyme, with CI4 as the predominant product. Therefore, substitution at residue F245 did not affect the strict product-size specificity of AgCI4Tase. Next, the effects of these mutations on CI4 formation and CI4 hydrolysis (a side reaction) were quantitatively measured as initial velocities (Table 2). The CI4 formation activity was reduced to less than one-fifth of the WT level in all mutants. In particular, F245W exhibited an approximately 180-fold reduction compared with the WT level. The hydrolytic activity of AgCI4Tase is weak, but was measurable under conditions of high substrate (10 mM CI4) and enzyme (1 μM) concentrations. The CI4 hydrolysis activity of F245A and F245L decreased to less than one-third of the WT level and was undetectable in F245W, whereas it increased more than twofold in F245Y. Taken together with the TLC results (Fig. 6), although cyclization reaction was reduced in F245A and F245L, the marked decrease in hydrolytic activity suppressed the degradation of produced CI4. Furthermore, reductions in disproportionation and coupling activities likely combined to suppress the formation of linear oligosaccharide by-products. In F245W, CI4 hydrolysis was almost abolished, but cyclization activity was also reduced, likely accounting for the reduction of disproportionation reaction products (IG2 and IG3). In F245Y, increased hydrolytic activity led to early the generation of IG4. Because its cyclization activity remained substantial, disproportionation and coupling reactions produced other linear oligosaccharides at levels comparable with WT.

**Fig. 7.**
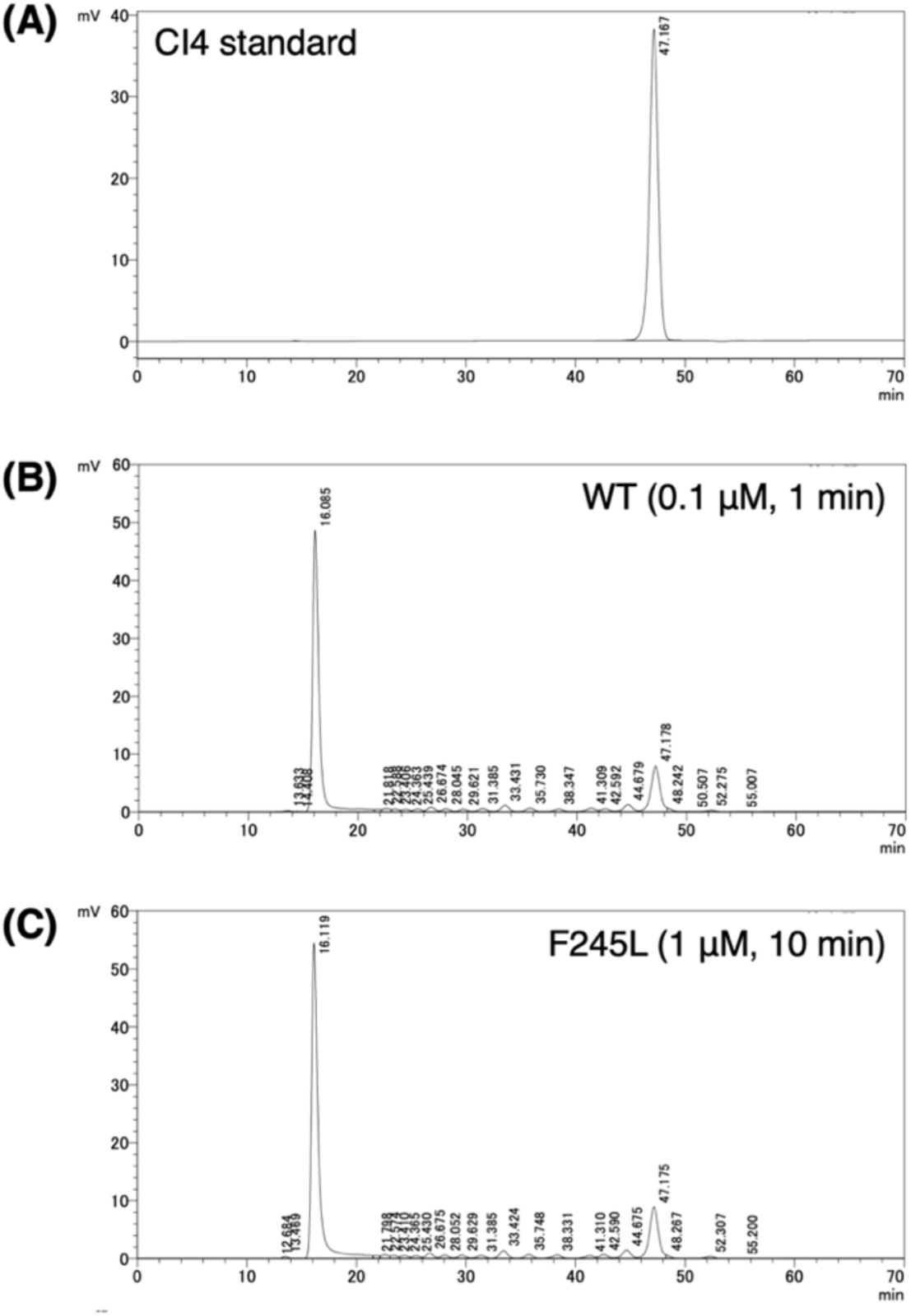
HPLC analysis of the product profile. (A) CI4 standard (10 mM). (B) Reaction products of WT AgCI4Tase. (C) Reaction products of F245L mutant. Reaction mixtures containing 1% (w/v) dextran, 50 mM Na-acetate (pH 6.0), and purified enzyme (indicated concentration) were incubated at 40°C for indicated time.

**Table 2.** Catalytic activities of wild type (WT) AgCI4Tase and F245 mutants.

|  | CI4 formation (min<sup>-1</sup>) | CI4 hydrolysis (min<sup>-1</sup>) |
| --- | --- | --- |
| WT | 285 ± 39 | 19.4 ± 2.0 |
| F245A | 15.6 ± 8.0 | 6.1 ± 1.2 |
| F245L | 29.6 ± 8.2 | 2.9 ± 0.7 |
| F245W | 1.6 ± 2.2 | N.D. |
| F245Y | 53.8 ± 15 | 58.6 ± 5.2 |

Initial velocities of the reactions at 40°C in 50 mM Na-acetate (pH 6.0) were measured. The CI4 formation (cyclization) from 1% dextran in the presence of 0.2 µM enzyme after 10 min was quantified using HPLC. The hydrolysis of 10 mM CI4 in the presence of 1 µM enzyme after 10 min was quantified by the production of glucose-equivalent reducing power using the BCA method. Note that the hydrolysis activity was measured at high concentrations of the cyclic substrate and enzyme, as the hydrolysis of CI4 formed by cyclization from dextran was barely detectable. See Materials and Methods for details. Values represent the mean ± standard deviation of three independent experiments. N.D. indicates that the activity was below the detection limit.

## Discussion

### Proposed mechanism for the strict production of cyclic tetrasaccharide

Based on our structural and kinetic analyses, we propose a structural model explaining the strict product specificity of AgCI4Tase (Fig. 8). The protruding loop (residues M247–R251) functions as a “molecular ruler” by creating a physical barrier that restricts the depth of the substrate-binding pocket (Fig. 5A). This steric hindrance ensures that only an isomaltooligosaccharide chain comprising exactly four glucose units is accommodated in the negative subsites, thereby defining the DP for subsequent cyclization. This structural feature of AgCI4Tase contrasts with that of conventional CITases, in which a spacious, elongated binding cleft, supported by the 2^nd^ site of CBM35 (Fig. 5B), allows long dextran chains to occupy multiple negative subsites (up to subsite – 8), facilitating the formation of larger macrocycles such as CI7 and CI8. In the cyclization reaction of AgCI4Tase, the O6 hydroxy of Glc(–4) performs an intramolecular nucleophilic attack on the anomeric C1 carbon of Glc(–1) (positioned approximately 16 Å away), resulting in the formation of the small cyclic product CI4 (Fig. 8). Within this constrained active site, residue F245 is positioned at the center of the bound cyclic product CI4 (Fig. 5C). Our mutational analysis demonstrated that F245 is essential for maintaining high cyclization activity in AgCI4Tase (Table 2).

**Fig. 8.**
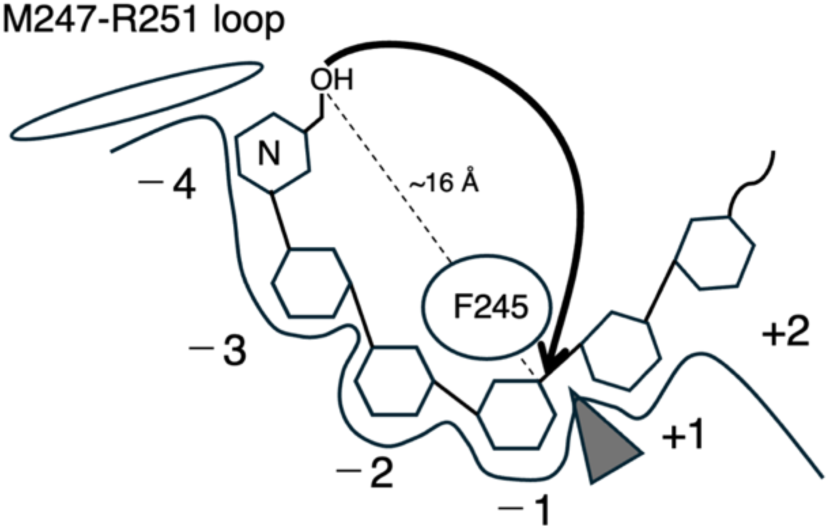
Proposed model for the strict four-glucose-unit specificity and reaction control in AgCI4Tase. The protruding M247–R251 loop creates a steric barrier that limits the substrate-binding pocket as a “molecular ruler”, allowing only IG4 glucose units to be accommodated for the subsequent cyclization. Residue F245 is located at the center of the cyclic product CI4 and is critical for the catalytic activity of AgCI4Tase.

The partitioning between cyclization (transglycosylation) and hydrolysis is fundamentally determined by the competition between a sugar hydroxy group and a water molecule for nucleophilic attack on the covalent intermediate. Therefore, efficient intramolecular transglycosylation requires mechanisms to physically exclude water molecules from the catalytic center or suppress their activation, such as by covering the active site with flexible loops [28,29] or introducing mutations at the +1 subsite [30,31]. In AgCI4Tase, the hydrophobic environment of subsite +1 (the transglycosylation acceptor site) formed by F245, L291, and V337 likely contributes to high cyclization and low hydrolytic activities (Fig. 4B). The increase in hydrolytic activity observed in F245Y (Table 2) underscores the importance of maintaining a strictly hydrophobic microenvironment at this position to prevent water-mediated side reactions. Although the F245A, F245L, and F245W mutants, in which the phenylalanine side chain was replaced with other hydrophobic residues, all exhibited reduced CI4 hydrolytic activity, their CI4-producing activity was impaired even more severely. These results indicate that an appropriately compact hydrophobic pocket is required to favor CI4-forming cyclization over competing hydrolysis, disproportionation, and coupling reactions.

### The functional role of CBM35

The GH66 family, to which CITases belong, is classified into three distinct subgroups based on their domain architecture and catalytic properties: (i) dextranases lacking CBM35 that exhibit only hydrolytic activity (EC 3.2.1.11), (ii) dextranases with CBM35-like domains showing low CITase activity, and (iii) CITases possessing CBM35 that favor CI production over hydrolysis [32]. This classification underscores the general importance of the CBM35 domain for efficient cyclization. In BcCITase, a unique binding site (2^nd^ site) is located adjacent to the catalytic center, which is characteristic of this CITase subgroup (Fig. 5B). The 2^nd^ site interacts extensively with the non-reducing end of the substrate, and mutations in this region are known to alter the product profile [33]. However, in AgCI4Tase, we observed relatively weak hydrogen-bonding interactions between the substrate and the CBM35 domain (Fig. 4A), indicating that the strict product specificity of AgCI4Tase is not primarily dictated by this domain.

In addition to the 2^nd^ site that assists the cyclization of CI, the CBM35 domains of BcCITase and PsCITase contain a canonical ligand-binding site (the 1^st^ site) located on the concave surface and distal from the active site [16,17]. The 1^st^ site is alternatively called the canonical site or variable loop site because the location of this site is conserved across all subfamilies (I-IV) of CBM35, and the structural variability of loop regions in the β-jellyroll fold dictates ligand specificity for galacto-, manno-, urono-, or gluco-sugars [34]. Because crystal structures of BcCITase and PsCITase in complex with IG8 or CI8 at the 1^st^ site have been determined, this site has been proposed to capture substrate polysaccharides such as dextran and guide them to the active site [16,17]. In the crystal structure of AgCI4Tase, no sugar ligand was bound to the 1^st^ site of the CBM35 domain, and only a glycerol molecule was observed (Fig. 5A, bottom). Comparison of the CBM35 domain of AgCI4Tase with those of BcCITase and PsCITase revealed that not only the central calcium-binding site, but also all residues involved in ligand binding are conserved (Fig. 9). Therefore, the 1^st^ site of the AgCI4Tase CBM35 domain is likely to have the same substrate-handling function as that of the CITases.

**Fig. 9.**
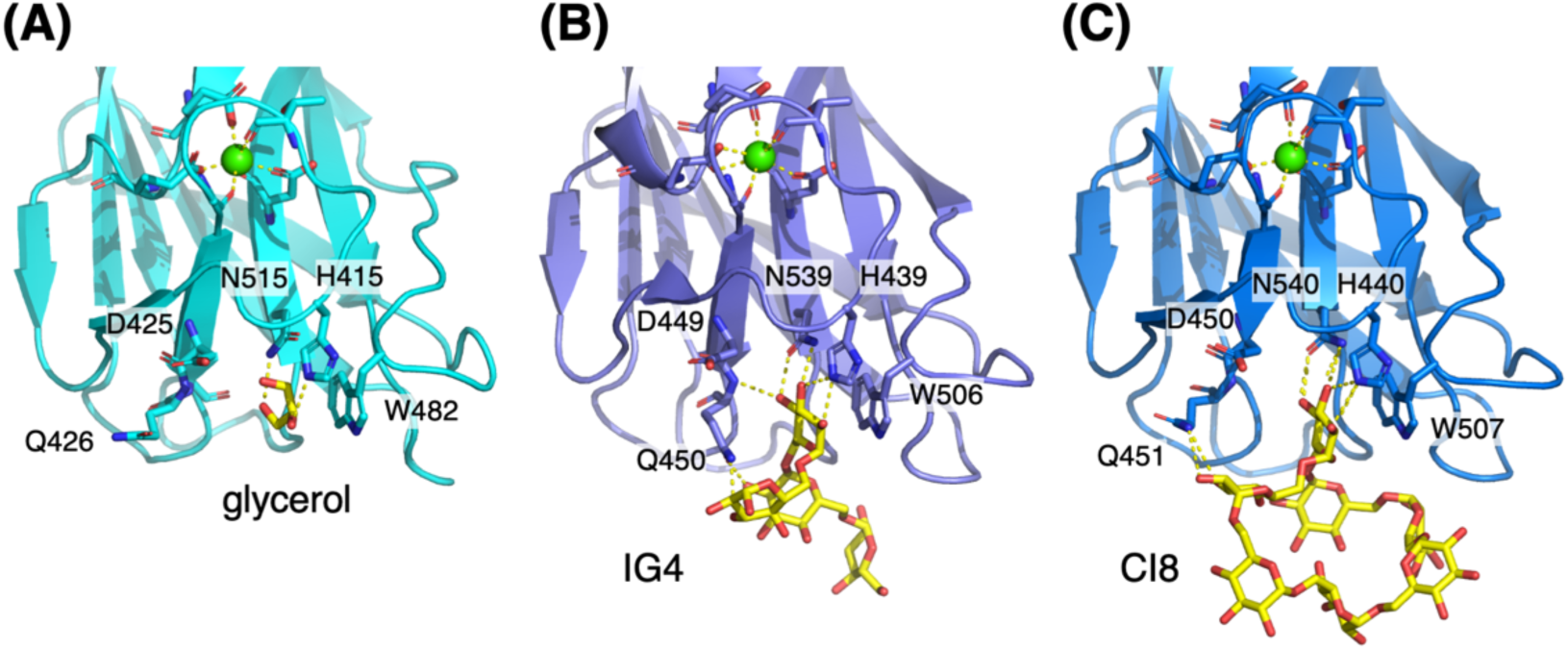
Comparison of the CBM35 domains of (A) AgCI4Tase complexed with glycerol, (B) BcCITase complexed with IG8 (PDB ID: 3WNN), and (C) PsCITase complexed with CI8 (PDB ID: 5X7H).

### Sequence landscape of GH66-related proteins and possible structural determinant for the functional diversity of CITases

To investigate the distribution of proteins homologous to AgCI4Tase, a sequence similarity network (SSN) analysis was performed using the top 2,994 sequences with the lowest *E*-values retrieved using BLAST (Fig. 10). This SSN covers a sequence space approximately fivefold larger than that of the 594 GH66 members currently listed in the CAZy database (https://www.cazy.org/GH66.html) and includes 21 characterized enzymes (CI4Tases, CITases, and dextranases). AgCI4Tase and MtCI4Tase (green diamonds) are located in the same subcluster as CITases (green circles). A dextranase with low CITase activity (PsDex) [32] is located in a small cluster (blue) adjacent to the CITase/CI4Tase cluster. A larger cluster containing only dextranases (yellow) is also located adjacent to the CITase/CI4Tase cluster, and these clusters were not separated at an alignment score cutoff of 100. The remaining bacterial dextranases are scattered among several relatively small, distinct clusters (cyan, red, and magenta). Several large clusters contain no functionally characterized enzymes, suggesting that they may harbor enzymes with novel activities.

**Fig. 10.**
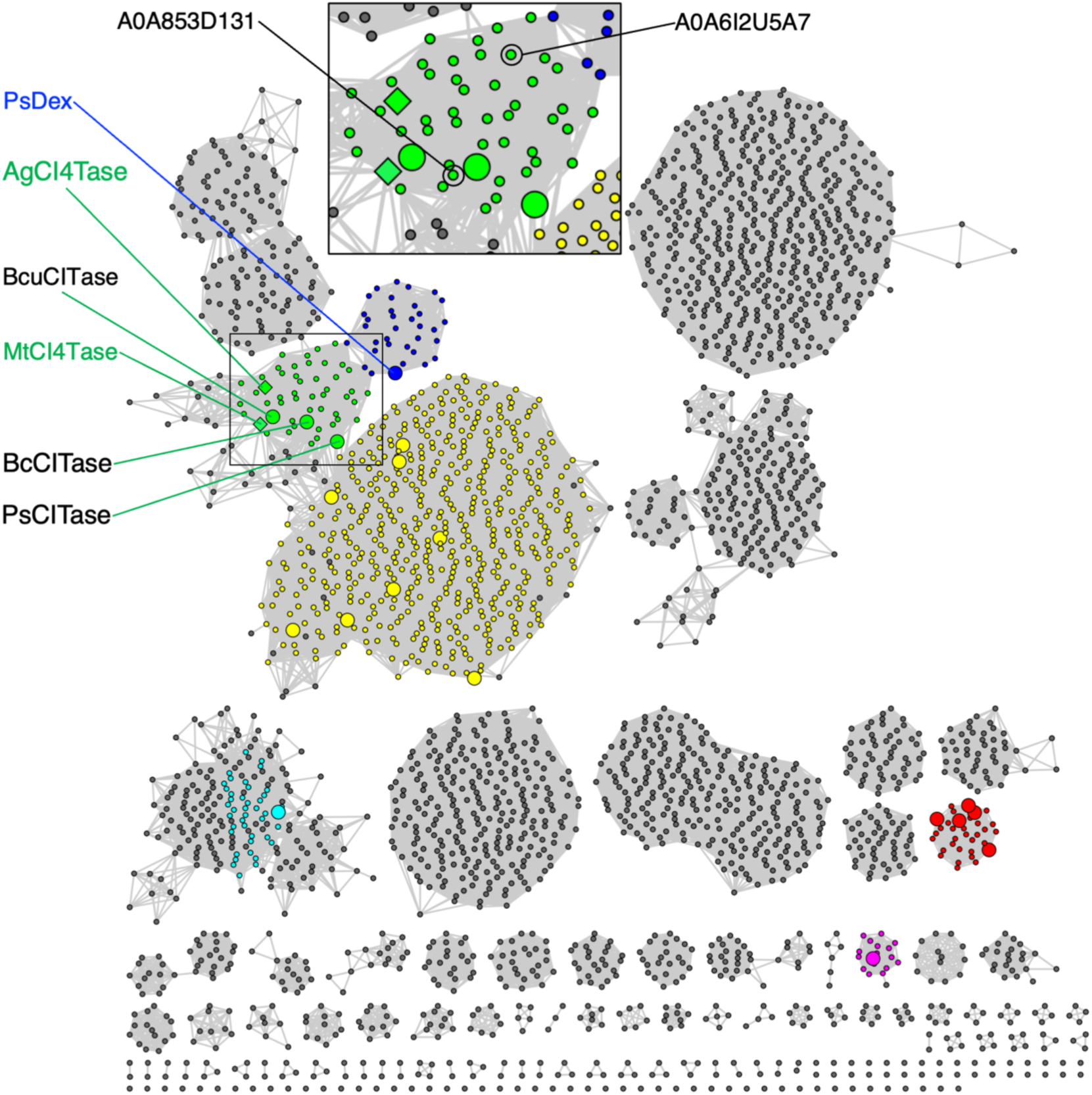
Sequence similarity network (SSN) of AgCI4Tase and homologous proteins. AgCI4Tase and MtCI4Tase are represented by green diamond nodes. Nodes representing characterized enzymes in the CAZy database (CITase, CI4Tase and dextranase) are enlarged. Green: BcCITase, PsCITase, and CITase from *Bacillus circulans* U-155 (BcuCITase) [49]. Nodes of the two putative proteins characterized in Fig. 12 are indicated in black circles (inset). Blue: Dextranase with low CITase activity from *Paenibacillus* sp. (PsDex) [32]. Yellow: dextranases from *Paenibacillus* spp. [50,51], *Thermoanaerobacter pseudethanolicus* [52], *Bacteroides thetaiotaomicron* [53], and *Flavobacterium johnsoniae* [54]. Cyan: dextranase from *Paenibacillus* sp. 598K [55]. Red: dextranases from *Streptococcus* spp. [56–59]. Magenta: dextranase from *Thermotoga lettingae* [60].

The intermixing of CITases and CI4Tases within a single, relatively small cluster in the SSN suggests that their product specificity is determined by local amino acid sequence differences rather than by overall sequence similarity. Focusing on the region corresponding to the “molecular ruler”, which our structural analysis identified as the key determinant of product specificity in AgCI4Tase, we generated a multiple sequence alignment of the proteins within the CITase/CI4Tase cluster (Fig. 11). This analysis revealed notable diversity in both the sequence and length of this loop region. Next, the crystal structures of AgCI4Tase, BcCITase, and PsCITase were superimposed on the AlphaFold-predicted structures of MtCI4Tase and two uncharacterized proteins (Fig. 12). MtCI4Tase and A0A853D131 possess loop sequences similar to the M247–R251 loop of AgCI4Tase, and the conserved proline residue is predicted to block Glc(–5) of IG8 in the same manner. These observations suggest that A0A853D131 likely also exhibits CI4Tase activity. In addition, although the corresponding loop sequence of A0A6I2U5A7 differs from that of AgCI4Tase (Fig. 11), a glutamate residue appears to form a similar steric barrier, suggesting that this protein may also possess CI4Tase activity. These two putative CI4Tases are distributed across different positions within the SSN cluster (Fig. 10). The structural diversity of this loop region further suggests that CITases with currently unknown product specificities, such as enzymes producing CI5 or CI6, might exist.

**Fig. 11.**
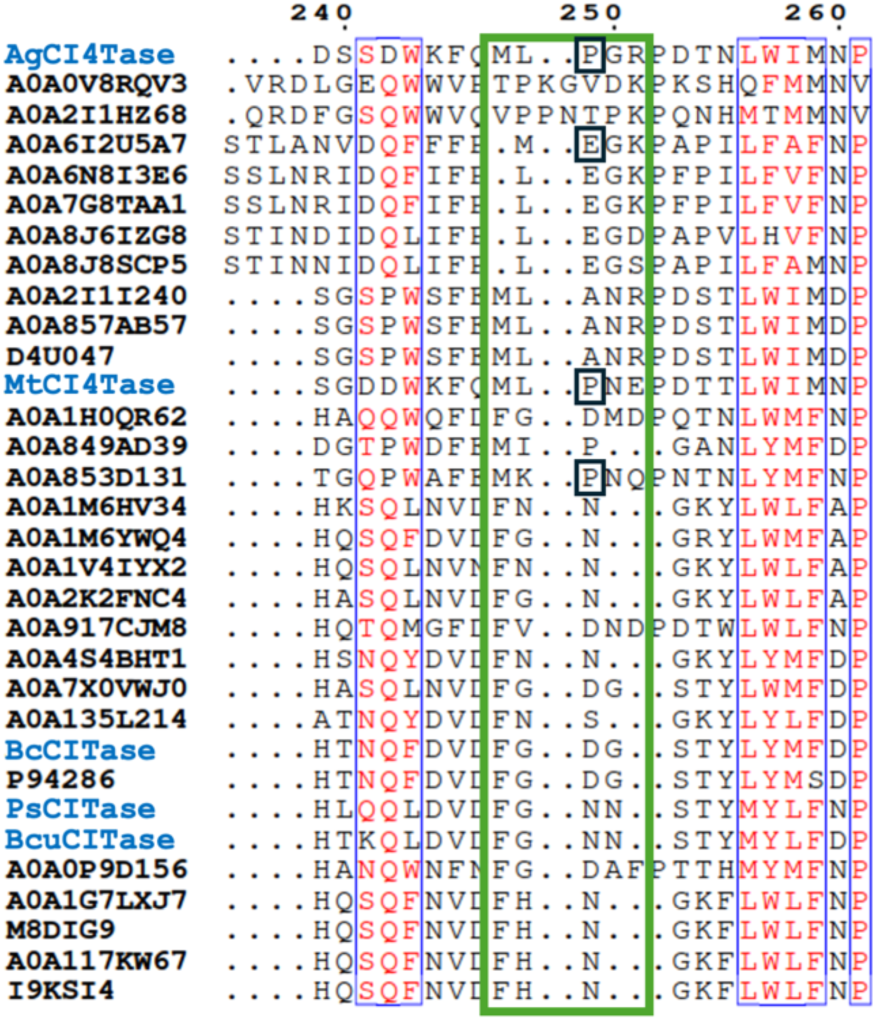
Multiple sequence alignment of the loop region in the CITase/CI4Tase cluster. The protruding loop region in AgCI4Tase (M247–R251) is enclosed by a thick green box. P249 in AgCI4Tase and the corresponding proline or glutamate residues in MtCI4Tase and two putative proteins, predicted to block subsite –5, are enclosed by black boxes.

**Fig. 12.**
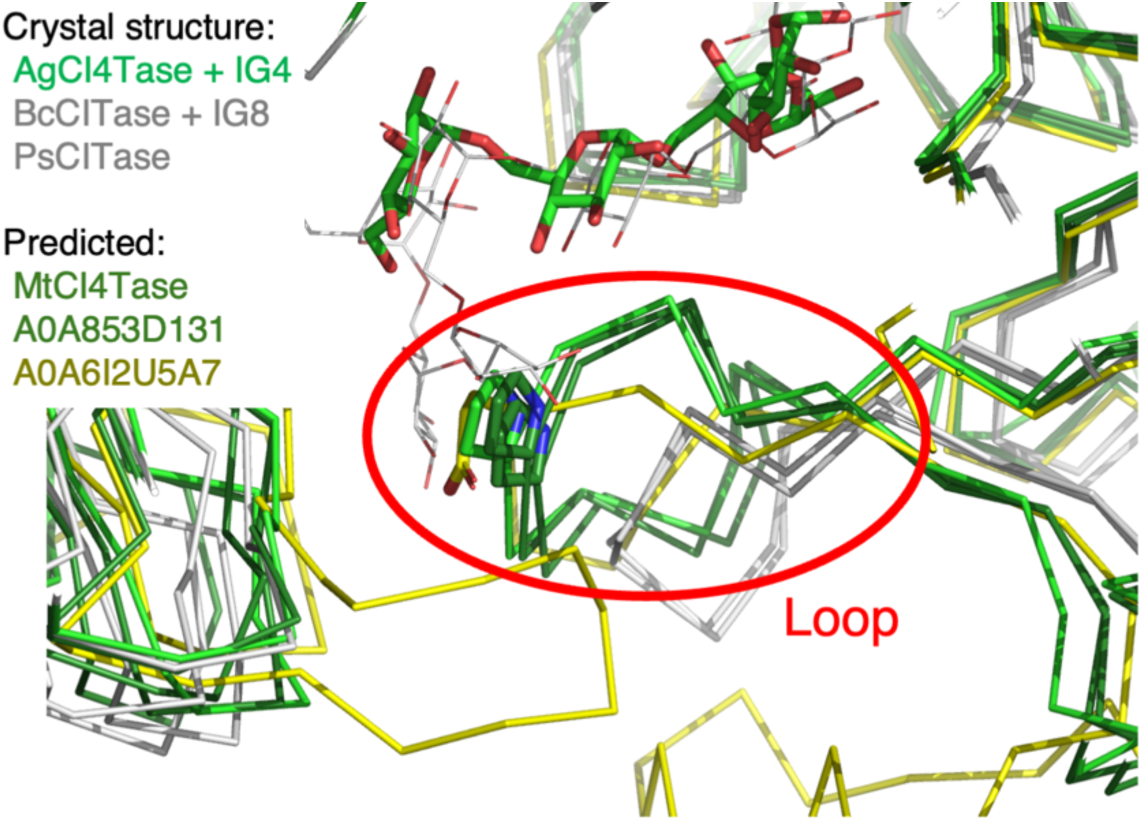
Predicted structural features of the protruding loops. The crystal structures of AgCI4Tase complexed with IG4 (green), BcCITase complexed with IG8 (gray), and PsCITase (gray) are superimposed with AlphaFold2-predicted structures of MtCI4Tase (dark green) and two putative proteins, A0A853D131 (dark green) and A0A6I2U5A7 (yellow). Only the Cα traces of the proteins and ligands in AgCI4Tase and BcCITase are shown.

## Conclusions

Obtaining enzymes that efficiently produce cyclic oligosaccharides with a defined DP while minimizing by-product formation is crucial for toward optimizing their cost-effective industrial production for diverse applications. In this study, we elucidated the “molecular ruler” mechanism underlying the strict product specificity of AgCI4Tase, which exclusively produces the novel cyclic tetrasaccharide CI4 via cyclization reaction. We further demonstrated that several F245 mutants retained the ability to produce CI4 while reducing the formation of linear IGs as by-products. In addition, our SSN analysis indicates that uncharacterized GH66-related proteins may harbor enzymes with novel activities. These findings provide an important structural framework for the rational design of transglycosylases tailored for specialized carbohydrate production.

## Materials and methods

### Protein production and purification

From the previously constructed expression plasmid pRSETA-ORF9038(-pep) [21], a region lacking the C-terminal CBM13 domain was amplified by PCR using the primer pair pET26-9038 (Table S1) and cloned into pET26b using the SLiCE method [35]. The SKIK sequence was subsequently introduced by PCR using the primer pair 9038 SKIK (Table S1), followed by DpnI (TaKaRa Bio Inc. Shiga, Japan) digestion and transformation into *Escherichia coli* DH5α. Expression plasmids for the D289A mutant and the F245 mutants were generated by site-directed mutagenesis using the primer pairs listed in Table S1 via PCR amplification and DpnI digestion. PCR amplification was performed using PrimeSTAR Max DNA Polymerase (TaKaRa Bio) or KOD One PCR Master Mix (TOYOBO Co. Ltd., Osaka, Japan), and PCR products were purified using the FastGene Gel/PCR Extraction Kit (NIPPON Genetics Co. Ltd., Tokyo, Japan). The plasmids were transformed into *E. coli* BL21 (DE3) CodonPlus-RIL cells. Cells were grown in Terrific broth medium containing 50 μg/mL kanamycin and 17 μg/mL chloramphenicol at 37°C until the OD600 reached 0.4 to 0.6. Protein expression was then performed at 20°C for 20 h without isopropyl β-D-thiogalactopyranoside induction. The harvested cells were resuspended in Buffer A (50 mM Tris-HCl pH 7.5 and 300 mM NaCl) and disrupted by sonication. After centrifugation (13,000×*g*, 45 min, 4°C), the supernatant was loaded onto a cOmplete His-Tag (Roche Diagnostic GmbH, Mannheim, Germany) column. The column was washed with Buffer A containing 5 mM imidazole, and the target protein was eluted with Buffer A containing 500 mM imidazole. The protein was further purified by size-exclusion chromatography using a Superose 6 10/300 GL column (Cytiva, Marlborough, MA, USA) equilibrated with 20 mM Tris-HCl pH 7.5 and 300 mM NaCl. The purity of the protein was confirmed by SDS-PAGE. The protein concentration was determined by measuring the absorbance at 280 nm using a NanoDrop spectrophotometer (Thermo Fisher Scientific, Waltham, MA, USA).

### Crystallography

The purified protein was concentrated and desalted before crystallization using Amicon Ultra Centrifugal Filter, 10 kDa MWCO (Merck, Darmstadt, Germany). Crystals were grown at 20°C using the sitting-drop vapor-diffusion method. A protein solution (0.5 μL) containing 15 mg/mL AgCI4Tase in 20 mM Tris-HCl (pH 7.5) was mixed with a reservoir solution (0.5 μL) containing 0.2 M NaCl, 0.1 M imidazole (pH 8.0), and 1.0 M K/Na tartrate. The IG4 complex was obtained by co-crystallization of the WT enzyme with 20 mM CI4. The CI4 complex was obtained by co-crystallization of the D289A mutant with 10 mM CI4. Crystals were cryoprotected by brief immersion (about 10 seconds) in a reservoir solution supplemented with 20% (v/v) ethylene glycol (ligand-free and CI4 complex) or 20% (v/v) glycerol (IG4 complex). The crystals were flash-cooled by dipping them into liquid nitrogen before data collection. X-ray diffraction data were collected at the beamlines of the Photon Factory of the High Energy Accelerator Research Organization (KEK, Tsukuba, Japan) and SPring-8 (Hyogo, Japan). The data were indexed, integrated, and scaled using XDS [36]. The structure of AgCI4Tase was determined by molecular replacement using PHASER [37] and a structural model predicted by ColabFold [38] as a search model. The model was refined using COOT [39] and Refmac5 [40]. Polder maps were prepared using the PHENIX software [41]. The final model was validated using MolProbity [42]. Molecular graphics images were prepared using PyMOL (Schrödinger LLC, New York, NY, USA).

### Enzyme assay

For TLC analysis, reaction mixtures containing 1% (w/v) dextran (average molecular weight 50,000–70,000) and 0.1 mg/mL enzyme in 50 mM Na-acetate (pH 6.0) were incubated at 37°C. Aliquots collected at 10, 30, 60, and 120 min, and after overnight incubation, were heat-inactivated at 100°C for 5 min. Samples (1 μL) were spotted onto Silica Gel 60 F_254_ sheets (Merck) and developed with a solvent system of 1-butanol/acetic acid/water (2:1:1, v/v/v). Carbohydrate spots were visualized by dipping the sheets in a diphenylamine-aniline-phosphoric acid staining reagent [43] followed by heating.

The cyclization activity was measured using HPLC. The reaction mixture (500 μL) consisted of 50 mM Na-acetate (pH 6.0) containing 1% (w/v) dextran. The reaction was initiated by adding the enzyme and the mixture was incubated at 40°C. For measurement of the product profile, 0.1 μM (WT) or 1 μM (mutant) enzyme was added (final concentration) and incubated for 1 min (WT) or 10 min (mutant), and then heat-inactivated at 100°C for 5 min. To measure the initial velocity of CI4 production, 0.2 μM enzyme was added (final concentration) and incubated for 10 min, and then the reaction was terminated by the addition of 75 μL of 100% (w/v) trichloroacetic acid. The mixture was incubated on ice for 30 min to ensure complete protein precipitation, followed by centrifugation at 13,000 rpm for 5 min. A 530 μL aliquot of the resulting supernatant was collected and neutralized with NaOH. The HPLC analysis was performed using a Prominence system (Shimadzu Corporation, Kyoto, Japan) equipped with a refractive index detector RID-20A. Samples (20 μL) were first filtered using a Millex-LG (0.2 μm; Merck) and desalted by electrodialysis using a micro acilyzer G0 (Asahi Chemical Co., Tokyo, Japan). Tandem MCI GEL CK04SS columns (10 mm i.d. × 200 mm × 2; Mitsubishi Chem. Co., Tokyo, Japan) were used at a flow rate of 0.4 mL/min using ultrapure water as a solvent at 80°C. Specific activity was expressed as min^-1^, calculated as the moles of cyclic product formed per mole of enzyme per minute. All measurements were performed independently three times.

The CI hydrolysis activity was measured by the bicinchoninic acid (BCA) assay [44]. The reaction mixture (100 μL) consisted of 50 mM Na-acetate (pH 6.0) and 10 mM CI4. The reaction was initiated by adding the enzyme to a final concentration of 1 μM and the mixture was incubated at 40°C for 10 min. The reaction was terminated by heat inactivation at 100°C for 5 minutes. For the blank, the enzyme solution was pre-inactivated at 100°C for 5 minutes before being mixed with the substrate. All reactions were performed in triplicate. The amount of reducing sugars was quantified using the TaKaRa BCA Protein Assay Kit (TaKaRa Bio). Working solution C was prepared by mixing Solution A (BCA solution) and Solution B (CuSO_4_ solution) in a 100:1 ratio. The reaction mixture was diluted with 50 mM sodium acetate buffer (pH 6.0) to fall within the detectable range.

Then, 75 μL of the diluted sample was mixed with 75 μL of Solution C in a 96-well plate. The plate was incubated at 70°C for 40 minutes, followed by stabilization at room temperature for 10 minutes. The absorbance at 562 nm was measured using a Synergy H1 microplate reader (Agilent Technologies Inc., Santa Clara, CA, USA). A standard curve was measured using D-glucose at concentrations of 0, 20, 40, 60, 80, and 100 μM. The mean absorbance of the blank was subtracted from each sample, and the glucose-equivalent concentration was calculated. The specific activity (min^-1^) was determined as the moles of reducing ends formed per mole of enzyme per minute. All measurements were performed independently three times.

### Sequence alignment and SSN analysis

Multiple amino acid sequence alignment was performed using the Clustal Omega [45] and ESPript [46]. A SSN was constructed using the EFI-Enzyme Similarity Tool (EFI-EST) server [47]. To identify homologous proteins for the network, the UniProtKB database was queried using the primary amino acid sequence of AgCI4Tase. The resulting network was visualized and analyzed using Cytoscape [48]. Clusters containing characterized CITases and CI4Tases were selected for further structural and sequence comparisons.

## Acknowledgments

We thank Mr. Takumi Fujita and Ms. Mayuko Matsumoto for their efforts in the preparation of the expression vector and initial crystallization screening, and Mr. Noriaki Kitagawa, Mr. Hiroki Asakuma, Dr. Masahiro Sota, and Mr. Takumi Masaki for their assistance in promoting collaborative research. We also thank Drs. Takatoshi Arakawa, Chihaya Yamada, Hajime Aga, Shimpei Ushio, and Koryu Yamamoto for their invaluable technical assistance and insightful discussions. We acknowledge the staff at SPring-8 and the Photon Factory for their support with X-ray data collection. This research was supported partly by Research Support Project for Life Science and Drug Discovery [Basis for Supporting Innovative Drug Discovery and Life Science Research (BINDS)] from AMED under the grant numbers: JP22ama121001, JP23ama121001, JP24ama121001, and JP25ama121001.

## Author contributions

SF, TM, and HW conceived and supervised the study. RY, TM, AM, and SF planned the experiments. RY, TM, and YK performed the experiments. RY and TK performed the protein crystallography. RY, TM, and YK performed the biochemical experiments. RY and SF wrote the manuscript. All authors reviewed the final version of the manuscript.

## Data availability statement

The nucleotide sequence of the AgCI4Tase gene has been deposited in the DDBJ/ENA/GenBank databases under accession number LC922692. The atomic coordinates and structure factors (PDB codes: 24TK, 24TL and 24TM) have been submitted to the Protein Data Bank (https://www.wwpdb.org/).

## Abbreviations

AgCI4Tase: *Agreia* sp. D1110 CI4Tase
BCA: bicinchoninic acid
BcCITase: Bacillus circulans CITase
CBM: carbohydrate-binding module
CD: cyclodextrin
CGTase: cyclodextrin glucanotransferase
CI: cycloisomaltooligosaccharide
CI4: cycloisomaltotetraose
CI4Tase: cycloisomaltotetraose glucanotransferase
CITase: cycloisomaltooligosaccharide glucanotransferase
CMM: cyclic maltosyl-maltose
CNN: cyclic nigerosylnigerose
DP: degree of polymerization
GH: glycoside hydrolase
HPLC: high-performance liquid chromatography
IG: isomaltooligosaccharide
IG4: isomaltotetraose
MtCI4Tase: *Microbacterium trichothecenolyticum* D2006 CI4Tase
PsCITase: *Paenibacillus* sp. 598K CITase
*R*_f_: retardation factor
RMSD: root-mean-square deviation
SSN: sequence similarity network
TLC: thin-layer chromatography
WT: wild type.

## Supporting Information

**Supplementary Table S1.**
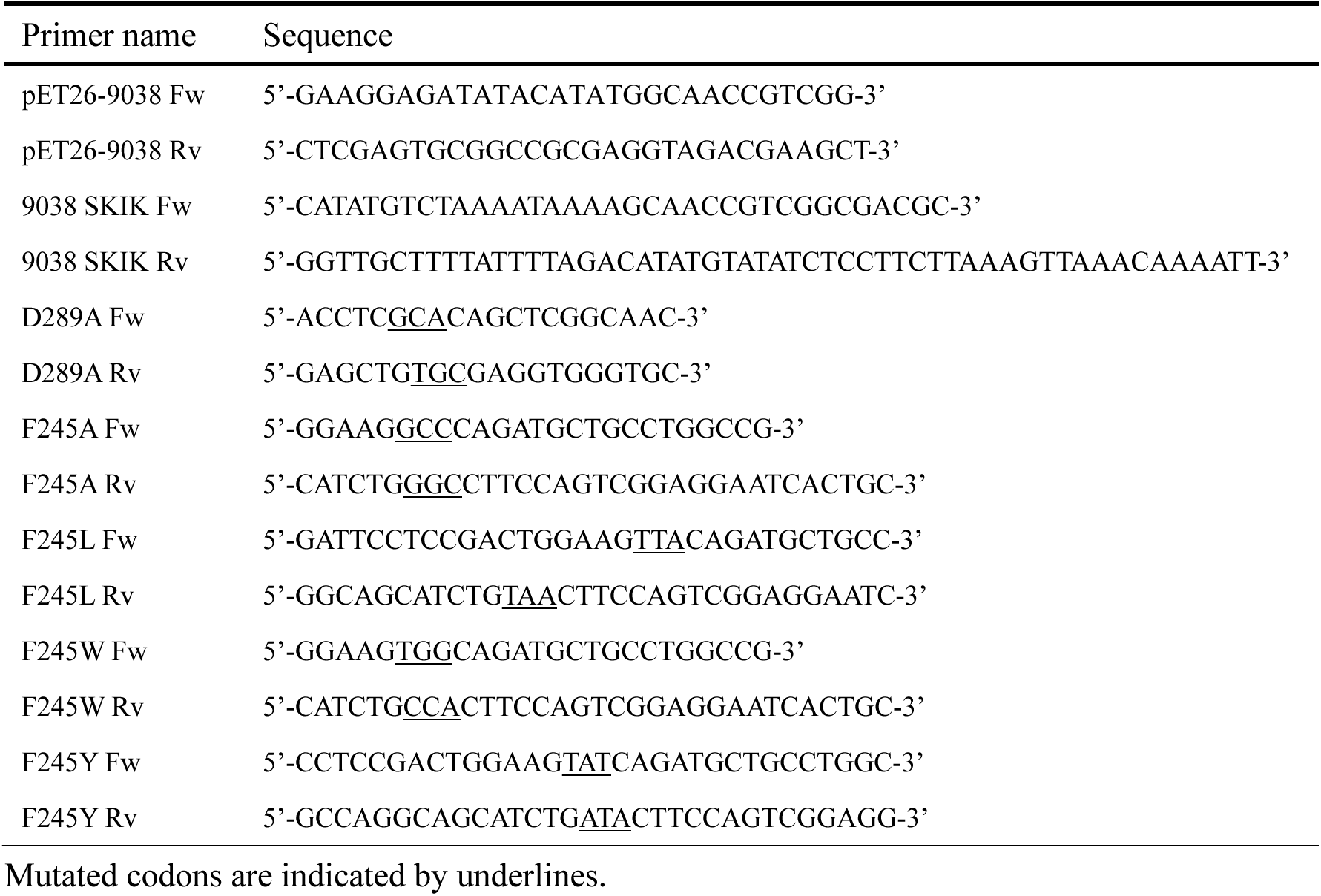
Primers used in this study Primer name Sequence.

